# The long and short of it: Benchmarking viromics using Illumina, Nanopore and PacBio sequencing technologies

**DOI:** 10.1101/2023.02.12.527533

**Authors:** Ryan Cook, Nathan Brown, Branko Rihtman, Slawomir Michniewski, Tamsin Redgwell, Martha Clokie, Dov J Stekel, Yin Chen, David J Scanlan, Jon L Hobman, Andrew Nelson, Michael A Jones, Darren Smith, Andrew Millard

**Author notes:** **Repositories**, All reads from virome sequencing were submitted to the ENA under the study PRJEB56639.

## Abstract

Viral metagenomics has fuelled a rapid change in our understanding of global viral diversity and ecology. Long-read sequencing and hybrid approaches that combine long and short read technologies are now being widely implemented in bacterial genomics and metagenomics. However, the use of long-read sequencing to investigate viral communities is still in its infancy. While Nanopore and PacBio technologies have been applied to viral metagenomics, it is not known to what extent different technologies will impact the reconstruction of the viral community.

Thus, we constructed a mock phage community of previously sequenced phage genomes and sequenced using Illumina, Nanopore, and PacBio sequencing technologies and tested a number of different assembly approaches. When using a single sequencing technology, Illumina assemblies were the best at recovering phage genomes. Nanopore- and PacBio-only assemblies performed poorly in comparison to Illumina in both genome recovery and error rates, which both varied with the assembler used. The best Nanopore assembly had errors that manifested as SNPs and INDELs at frequencies ~4x and 120x higher than found in Illumina only assemblies respectively. While the best PacBio assemblies had SNPs at frequencies ~3.5 x and 12x higher than found in Illumina only assemblies respectively. Despite high read coverage, long-read only assemblies failed to recover a complete genome for any of the 15 phage, down sampling of reads did increase the proportion of a genome that could be assembled into a single contig.

Overall the best approach was assembly by a combination of Illumina and Nanopore reads, which reduced error rates to levels comparable with short read only assemblies. When using a single technology, Illumina only was the best approach. The differences in genome recovery and error rates between technology and assembler had downstream impacts on gene prediction, viral prediction, and subsequent estimates of diversity within a sample. These findings will provide a starting point for others in the choice of reads and assembly algorithms for the analysis of viromes.

**Data Summary:** All reads from virome sequencing were submitted to the ENA under study PRJEB56639. The assemblies are provided via FigShare (https://figshare.com/s/2d9b5121eb421d370455).

**Author Notes:** Eight Supplementary Tables and nine Supplementary Figures are available with the online version of this article.

## Introduction

Viruses play critical roles in all environments they inhabit, as evidenced by their distribution and abundance. In particular viruses that infect bacteria, bacteriophages (from hereon phages), are known to play important roles in regulating the abundance of their bacterial hosts, facilitating horizontal gene transfer and playing crucial roles in global biogeochemical cycles by augmenting host metabolism [1–3].

It is now over 40 years since sequencing of the first phage genome [4]. The number of complete phage genomes from phage isolates is now >22,000 [5]. However, millions more phage genomes have been sequenced through metagenomics sequencing and are available through a variety of databases [6–8]. Viral metagenomics (viromics) has revolutionised our understanding of the diversity of phages and their potential ability to augment host metabolism. Initial virome studies required DNA to be cloned into a vector and the clone sequenced by Sanger sequencing. As new sequencing technologies developed that did not require the cloning of DNA, such as Solexa (becoming Illumina), 454 and SOLiD, the field of viromics expanded. With Illumina sequencing becoming the dominant technology, more and more viromes have been sequenced spanning pristine ocean environments [9], abyssal depths [10], and even the faeces of a wide variety of animal species [11–13].

Whilst viromes produced using Illumina short-read sequencing have provided new insights into viral diversity, short reads are not able to resolve all viral genomes within a virome. Phages that contain hypervariable regions and/or possess high microdiversity are known to cause virome assemblies to fragment, resulting in reduced contig sizes and exclusion from further analyses [14]. To overcome such problems, alternative approaches to viromics can be taken including single cell viromics or the cloning of viral genomes into fosmids [15]. Whilst both of these approaches are beneficial, they are technologically challenging compared to more standard viromics workflows.

Recent technological developments have led to the production of long read sequences by both Oxford Nanopore Technology (ONT) [16] and PacBio [17]. While the technologies differ in their approach, both platforms sequence single molecules and are capable of producing sequences of tens of kilobases in length [17]. The ability to sequence long DNA molecules offers the ability to overcome the issues of microdiversity and or hypervariable regions found within phage genomes [14]. To date there have been limited studies using ONT sequencing for viromics. One of the first was able to acquire complete phage genomes from single ONT reads, utilising tangential flow filtration (TFF) of marine samples to obtain the significant amounts of DNA required for library preparation [18]. Extraction of such quantities of phage DNA is likely prohibitive from more viscous and heterogeneous environments where multiple displacement amplification (MDA) is already used to obtain enough DNA for library preparation for short read sequencing. While MDA provides a solution to the amount of input material, it does not come without problems. It has been well documented that MDA can introduce biases in metagenomic libraries, in particular over representation of ssDNA phages within samples [19–21]. To overcome the problem of library input requirements, MDA for ONT library preparation, combined with unamplified short read libraries for quantification have been utilised [22]. Alternatively, ONT sequencing (minION) of long-read linker amplified shotgun libraries (LASL), to sequence PCR products on a minION, combined with Illumina short reads were used in an approach dubbed virION [14, 23]. Both approaches were successful in increasing the number and completeness of viral genomes.

While the number of viromes that utilise ONT alone or in combination with Illumina sequencing is slowly increasing [14, 22–25], reports of utilising PacBio sequencing for viromes are scarce [26]. A recent study predicted phages from a bacterial metagenome assembled from PacBio reads, identifying phages not identified when the same sample was sequenced with short reads [26]. It is not clear why there are not more viromes sequenced with long read technologies, as has become commonplace for sequencing of bacterial metagenomes. Even for the sequencing of individual phage isolates there are relatively few studies that have utilised long reads [27–30]. In part, this is likely because the vast majority of phage genomes can be assembled from short read Illumina sequences alone [31]. Thus, unlike sequencing their bacterial hosts, long reads do not provide the immediate benefit of a better genome assembly for an isolate and thus the need to use them is reduced. The lack of long-read data generally for phage isolates, combined with the lack of a comparative benchmarked dataset comparing different methods is likely contributing to long read sequencing not being widely adopted for viromes, despite clear benefits from the limited studies performed to date. We aimed to understand how different sequencing technologies affect the recovery of viral genomes from communities and the quality of the assembled genomes.

Here then, we sequenced a mock community of phages with three different sequencing technologies (PacBio, minION and Illumina) in order to benchmark these different approaches and identify the benefits and limitations of each approach.

## Methods

### Mock Virome Community Preparation and Sequencing

Phages (vB_Eco_SLUR29, vB_EcoS_swan01 [32], vB_Eco_mar001J1 [32], vB_Eco_mar002J2 [32], KUW1 (OQ376857), PARMAL1 (OQ376857), HP1 [31], DSS3_PM1, vB_Eco_mar005P1 [32], S-RSM4 [33], vB_Eco_mar003J3 [32], vB_Vpa_sm033, vB_VpaS_sm032, CDMH1 [31]) were propagated as previously described and DNA was extracted using the method of Rihtman et al (2016). DNA was quantified with the Qubit dsDNA high sensitivity kit. ΦX174 DNA was obtained from the spike in control provided with Illumina library preparation kits. Genomic DNA was combined to produce a mock community of fifteen phages that covered a range of lengths (44,509 - 320,253 bp) and molGC content (38% - 61%). Genomes were combined across a range of abundances (169,000 - 684,329,545 genome copies) within the mock community (Supplementary Table 1). Genome copies were estimated by using the formula: (ng of DNA * 6.022 x 10^23^) / (Genome Length * 660 * 1 x 10^9^). The genomes were chosen to include both highly divergent and highly similar phages (Supplementary Table 2; Supplementary Figure 1).

Illumina library preparation was carried out using the NexteraXT library preparation kit, with a minor modification to the number of PCR cycles as described previously [32]. In addition, no ΦX174 spike was added to the library as it is part of the normal Illumina library preparation protocol. Sequencing was carried out with a MiSeq 2 x 250 bp kit. For minION and PacBio sequencing, the DNA was amplified prior to sequencing with the GenomiPhi V3 DNA Amplification Kit, following the manufacturer’s instructions. Eight individual amplification reactions were performed with 10 ng DNA input for each amplification. Following amplification, DNA was treated with S1 nuclease with 10 U per μg of input DNA and the enzyme deactivated, prior to cleanup and concentration with a DNA Clean & Concentrator-25 column (Zymo Research). Three independent amplification reactions were sequenced via PacBio or ONT sequencing.

Libraries were prepared for minION sequencing using SQK-LSK109 (Version: NBE_9065_v109_revB_23May2018) with the native barcoding kit, following the manufacturer’s instructions (Oxford Nanopore Technologies, Oxford, UK) with omission of the initial g-tube fragmentation step. Base calling was carried out with Guppy v2.3.5, with reads demultiplexed using Porechop [34]. PacBio sequencing was carried out at NU-OMICS (Northumbria University). Briefly, genomic DNA was sheared using g-TUBE (Covaris, USA) to an average size of 8-10 kb and then libraries were prepared using SMRTbell Template Prep Kit 1.0 (Pacific BioSciences, USA) as per manufacturer’s instructions and sequenced on the Sequel I system (Pacific BioSciences, USA). Circular consensus reads were created in SMRTLink v6 (Pacific BioSciences, USA) and fastq files were generated using the BAM to FASTX pipeline.

### Bioinformatics Analyses

To determine coverage and depth, reads from each library were mapped to the 15 reference genomes using Minimap2 v2.14-r892-dirty with “-ax sr”, “-ax map-ont”, or “-ax map-pb” for Illumina, ONT and PacBio reads respectively [35]. Minimap2 output was piped and sorted using the Samtools sort command to produce sorted bam files [36]. Coverage and depth were taken from the bam files using the Samtools coverage command [36].

Assemblies were separately produced for the three libraries, and additional assemblies were produced by pooling the three libraries together, resulting in four assemblies per read/assembler combination. Illumina reads were trimmed with Trim Galore v0.4.3 prior to assembly [37]. Illumina reads were assembled using SPAdes v3.12.0 with parameters “--meta -t 16” [38]. Flye assemblies were produced with parameters “--nano-raw/or --pacbio-raw --threads 90 --meta” [39]. Unicycler assembly of long reads was used with default parameters [40], that utilise miniasm [41] for an overlay consensus assembly followed by racon for polishing [42]. wtdbg2 was used with the parameters “-p 21 -k 0 -AS 4 -K 0.05 -s 0.05 -L 1000 --edge-min 2 --rescue-low-cov-edges -t 90” [43].

To determine whether using long and short reads together improved the assemblies, three methods that utilised a hybrid approach were used. (1) Long read-only assemblies were polished with multiple rounds of polishing using Pilon [44] (hereafter referred to as “polished”). (2) For a hybrid assembly with Unicycler, long and short reads were provided with default parameters (hereafter referred to as “hybrid”). (3) The hybrid Unicycler assemblies were combined with the Illumina-only assemblies and de-replicated at 95% average nucleotide identity (ANI) over 80% genome length using the ClusterGenomes script [45].

To assess completeness and quality, assemblies were compared to the 15 reference genomes using metaQUAST v5.0.2 with default parameters [46]. All resultant plots were produced using ggplot2 in R v3.5.1. When investigating the fidelity of assemblies to the reference genomes, we included assemblies for which 50% of the genome was covered by contigs, no matter how fragmented the assembly was (i.e., if 100 individual contigs mapped to 50% of genome length, despite the longest contig only being 10% of genome length, this was still included. This was to exclude misassembly data for which only small portions of genomes were assembled, potentially leading to under-estimation of error frequencies). To investigate the effect of sequence depth on long read assembly, reads mapping to the genome of interest were extracted and downsampled using seqtk sample with -s100 to the desired depth.

To determine the effect of polishing long-read assemblies with short-reads on viral prediction software, we processed the long-read assemblies and their polished counterparts using VIBRANT v1.2.1 [47] with the following parameters “-t 8 -l 10000 -virome” and compared against DeepVirFinder v1.0 [48] with contigs >10 kb and a P-value <0.05. Prodigal v2.6.2 with default settings was used for predicting open reading frames on the vOTUs and the 15 reference genomes [49].

To investigate the effect of different sequencing platforms and assemblers on estimates of viral diversity, we applied a typical virome analysis workflow to the assemblies. Each assembly was separately processed using DeepVirFinder v1.0 [48]. Contigs ≥10 kb or circular were included as viral operational taxonomic units (vOTUs) if they obtained a P-value of <0.05. Reads from the corresponding Illumina library were mapped to the assembly using Bbmap v38.69 at 90% minimum ID and the ambiguous=all flag [50]. vOTUs were deemed as present in a sample if they obtained ≥ 1x coverage across ≥ 75% of contig length [51]. The number of reads mapped to present vOTUs were normalised to reads mapped per million. Relative abundance values were analysed using Phyloseq v1.26.1 [52] in R v3.5.1 to calculate diversity statistics [53]. The number of predicted vOTUs and alpha diversity statistics were compared to the genome copy numbers used in the original mock community.

## Results

### Mock Virome Composition

To assess the performance of short, long, and hybrid sequencing approaches for viromic analyses, we sequenced a mock community of 15 bacteriophage genomes with an Illumina MiSeq, PacBio Sequel, and ONT minION. For Illumina sequencing, no MDA was used to provide a library as free as possible from bias. For PacBio and ONT sequencing, the mock community was first amplified with MDA to obtain sufficient material for library preparation and sequencing. The Illumina and ONT libraries yielded similar amounts of data with 0.5 - 1.1 Gb and 0.6 - 1.1 Gb respectively, and 0.3 - 0.5 Gb from PacBio libraries. Pooling the libraries resulted in 2.4, 2.7 and 1.1 Gb for Illumina, ONT and PacBio libraries respectively (Supplementary Table 3).

### Limits of Detection by Read Mapping

First, we assessed the limits of detection of each sequencing platform using a mapping-based approach, with detection of a genome set at 1x coverage across ≥97% of a genome. Four phage genomes were not detected at all (CDMH1, HP1, vB_Eco_mar005P1 and ΦX174) by any sequencing technology (Figure 1A). The Illumina and ONT libraries detected a similar number of genomes (8-10 genomes), with PacBio detecting 7-8 genomes (Figure 1B). Although Illumina and ONT both recovered 8-10 genomes across all libraries, the average number of genomes detected in a single Illumina library was higher than that of a single ONT library (Figure 1B). The least abundant phage to be detected was S-RSM4 (465,530 copies) and was only detected by Illumina sequencing, although a small percentage of the genome was covered in the PacBio and ONT libraries. The least abundant phages detected in ONT and PacBio libraries were vB_VpaS_sm032 (52,465,265 copies) and J1 (53,672,906 copies), respectively.

**Figure 1.**
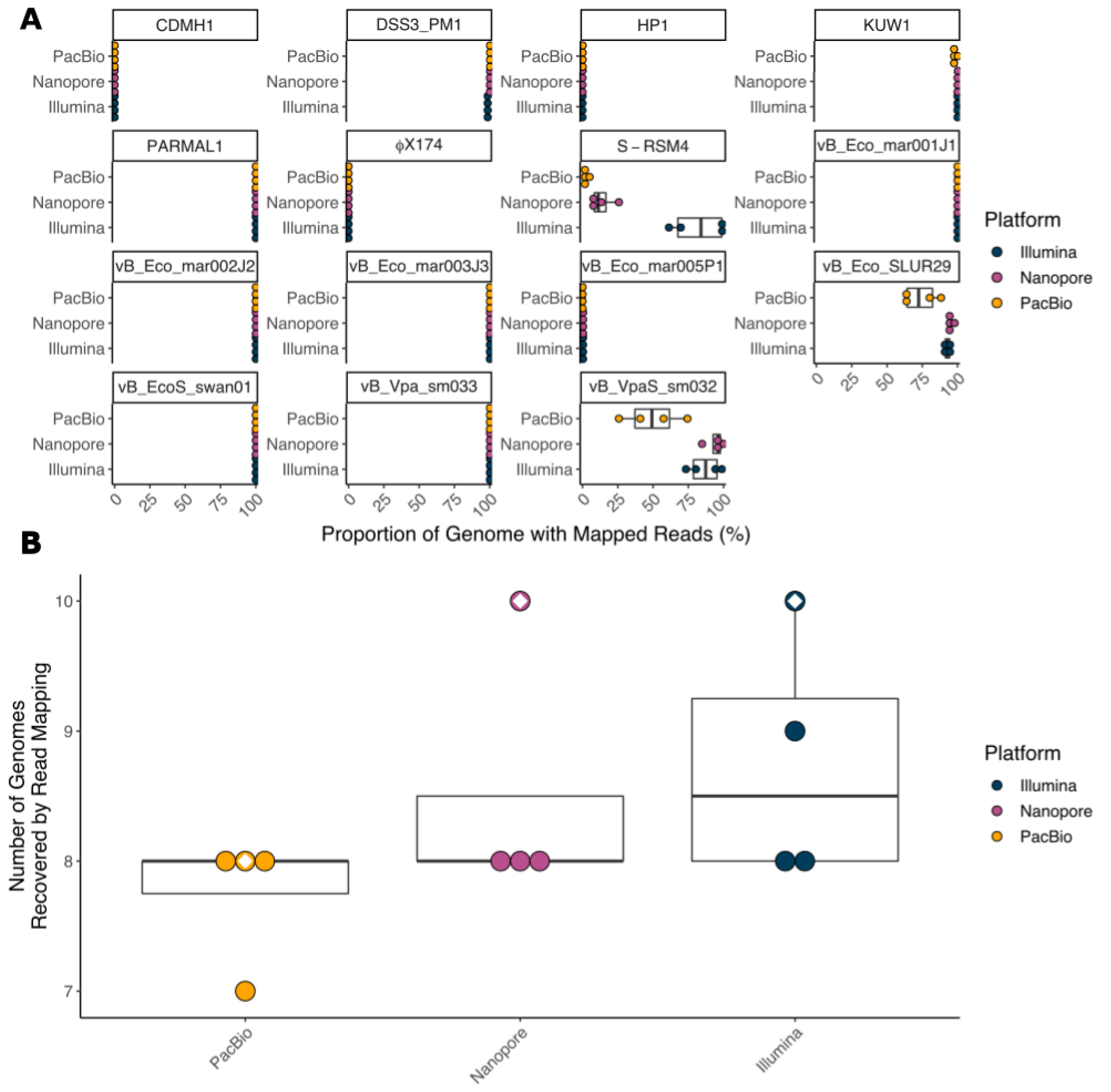
Detection of genomes by read mapping. (**A**) Dot and boxplots showing the proportion of each genome to which reads were mapped from each of the three sequencing platforms for library repeats, and (**B**) the number of genomes detected by read mapping by each sequencing platform at 1x coverage over ≥97% of genome length with the white diamond indicating the pooled library.

The use of unamplified DNA for Illumina libraries allowed any effects of MDA to be identified in the long read assemblies. Encouragingly, the abundance of a genome within a sample generally correlated across different sequencing platforms, even after MDA for PacBio and ONT sequencing (ONT vs Illumina r=0.9903948, PacBio vs Illumina r=0.9883086, ONT vs PacBio r=0.9996938) (Figure 2A, B, C; Supplementary Table 4). However, it should be noted that phage ΦX174 was not detected in any sample, suggesting we may have been overly cautious in the amount we added to the mock community.

**Figure 2.**
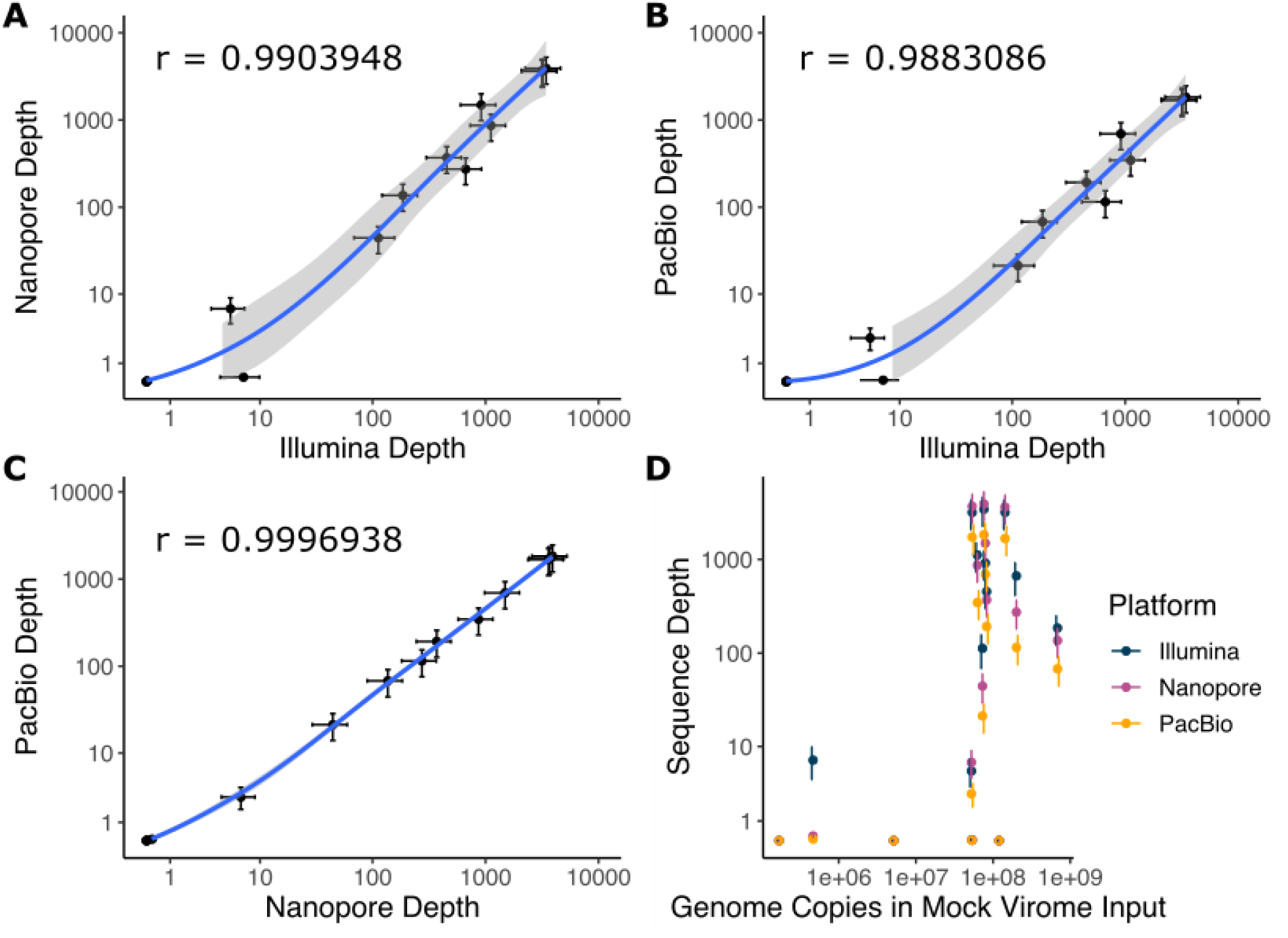
Comparison of sequencing depth between platforms. Correlation plots showing average sequence depth of a genome between (**A**) Illumina and ONT, (**B**) Illumina and PacBio, and (**C**) ONT and PacBio. An additional plot (**D**) shows sequence depth for the three sequencing platforms versus the estimated number of genome copies in the original mock community from which DNA libraries were prepared and sequenced. Values shown are the mean across three libraries and a pooled library, with bars showing standard error.

### Assembly Results - Genome Recovery

As assembly options for each read type were tested to optimise assembly methods, assemblies were obtained for all samples and assemblers tested, with the exception of PacBio reads using Unicycler (miniasm + racon) so these were excluded from further analysis. To investigate whether combining read technologies led to more complete assemblies, PacBio and ONT reads were separately assembled alongside Illumina reads using Unicycler to produce “hybrid” assemblies. The hybrid assemblies were separately combined with Illumina only assemblies and de-replicated at 95% average nucleotide identity (ANI) over 80% length to produce “deduped” assemblies [54].

For individual sequencing platforms, only short reads (Illumina) resulted in any completely assembled genomes (3-4) (Figure 3; Supplementary Figure 2). Despite having >1,000x coverage of some genomes in long-read-only libraries the reads did not assemble into complete genomes, suggesting that coverage is not a limitation and may well be a hindrance to assembly (Supplementary Table 4). The Illumina + ONT hybrid assembly (Unicycler) recovered the most genomes (2-6 genomes) (Figure 3; Supplementary Figure 2). The addition of long reads to short reads increases the number of genomes recovered (particularly ONT).

**Figure 3.**
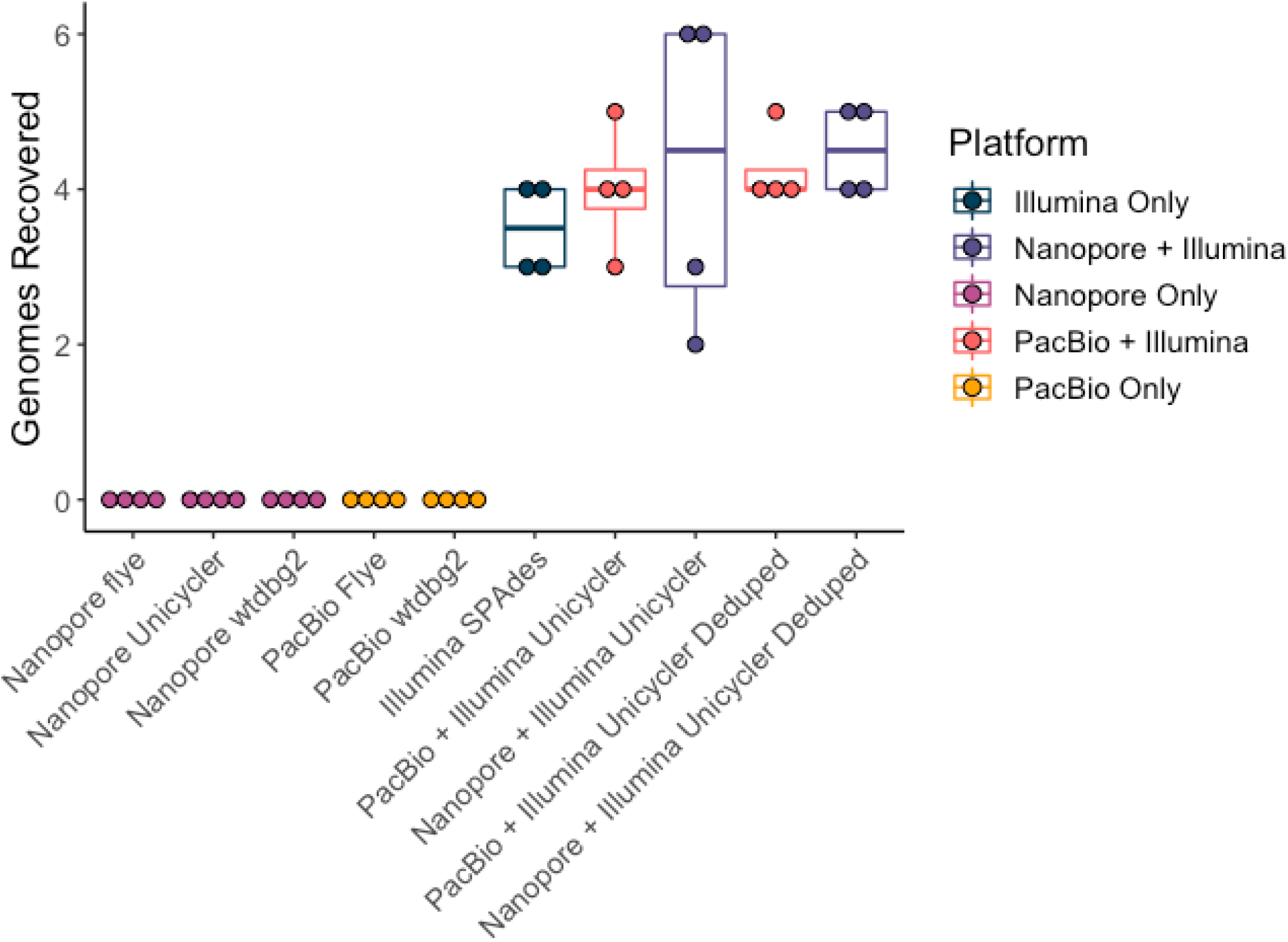
Comparison of genome recovery across sequencing technologies and assemblies. Dot and boxplot showing the number of genomes fully assembled within each assembly (successful assembly defined as a single contig covering 97% of genome length), with the reads used for assembly shown in different colours.

### Assembly Results - Limits of Detection for Assembled Genomes

The phage with the lowest input abundance to be recovered in a single contig within any assembly was vB_Eco_mar001J1 (53,672,905 genome copies), which was recovered in the ONT + Illumina hybrid assembly. The least abundant genome to be recovered from an Illumina only assembly, and a PacBio + Illumina hybrid assembly was KUW1 (72,995,151 genome copies), suggesting the addition of ONT reads to Illumina reads improves the recovery of lowly abundant genomes, but the addition of PacBio reads to the same Illumina reads did not.

Whilst not the genome with lowest input abundance, KUW1 was recovered from Illumina libraries at a lower average sequence depth than any other genome (139 x coverage in the largest Illumina library, 225 x coverage in the pooled library). Furthermore, KUW1 was not assembled in the two smaller Illumina libraries (37 and 49 x coverage obtained), suggesting that the depth of Illumina sequencing impacts the limits of detection.

As previously discussed (**Limits of Detection by Read Mapping**), the least abundant genomes to be detected by read mapping were vB_VpaS_sm032 and S-RSM4. Summed Illumina contigs from the pooled library mapped to 87% and 97% of vB_VpaS_sm032 and S-RSM4 respectively. However, the longest individual contigs only covered a small fraction of the genomes (22% and 9% respectively). The average read depth for vB_VpaS_sm032 and S-RSM4 contigs was 10 x over 98.7% and 14 x over 99.6% of genome lengths respectively in the pooled Illumina library. Manual inspection of alignments revealed that breaks in the assemblies typically coincided with a drop in read coverage which was often associated with a sudden and sharp change in mol%GC (either upwards or downwards; Supplementary Figure 3; Supplementary Table 4) [55–57].

The longest genome to be recovered in a single contig, vB_Vpa_sm033 (320,253 bp), was assembled in Illumina-only, Illumina + PacBio, and Illumina + ONT assemblies. The shortest genome to be recovered in a single contig, KUW1 (44,509 bp), was assembled in Illumina-only, Illumina + PacBio, and Illumina + ONT assemblies. Whilst KUW1 was assembled from only one individual Illumina library, it was assembled in two each of the ONT + Illumina, and PacBio + Illumina hybrid assemblies. Furthermore, dereplicating these hybrid assemblies with the Illumina-only assemblies led to KUW1 being assembled in all individual libraries.

### Assembly Results - Resolution of Highly Similar Genomes

It was possible to assemble a single genome that was representative of vB_Eco_mar001J1 and/or vB_Eco_mar002J2 from Illumina + ONT hybrid assemblies, rather than two genomes, perhaps unsurprisingly given there is >99% ANI between them (Supplementary Figure 1). A single genome of vB_EcoS_swan01 was obtained using an Illumina + ONT hybrid assembly, which has ~80% ANI with vB_Eco_SLUR29 (Supplementary Figure 1). However, the genome of vB_Eco_SLUR29 could not be resolved.

### Assembly Results - Comparison of Long Read Assemblers

Despite high read coverage, long-read only assemblies failed to recover a complete genome for any of the 15 phage (Figure 3; Supplementary Table 4). To identify the optimal long-read only assemblies, we used the NGA50 statistic (Supplementary Figure 4 & 5). While nine genomes were detected by mapping long reads in at least one library, only eight are included in this analysis due to the very low coverage of vB_Eco_SLUR29 recovered from any assembly. For this comparison, we also included long read assemblies that were polished with Illumina reads, as this was found to affect the results.

The NGA50 values averaged across the eight genomes and four libraries obtained from ONT assemblies were higher than those from PacBio. Again this varied depending on the assembler used. ONT reads assembled with Flye, wtdbg2 and Unicycler obtained average NGA50 values of 28%, 10% and 8% respectively, whereas PacBio reads obtained values of 5% and 4% for wtdbg2 and Flye assemblies respectively. ONT reads assembled with Flye typically produced the longest alignments in relation to reference genomes. The performance of Flye with individual libraries was higher than that in the pooled library, with the average NGA50 values as a proportion of genome length being 27%, 39% and 30% for individual libraries, and only 16% for the pooled library (Supplementary Figure 4 & 5). Conversely, the highest NGA50 values for ONT reads assembled with Unicycler were obtained from the pooled library (26%), and 3%, 1% and 0.1% from individual libraries (Supplementary Figure 4 & 5). Thus, pooling reads was both detrimental or advantageous, depending on the assembler used.

For all five long-read only assemblies, its polished counterpart typically obtained more complete assemblies than before polishing (Supplementary Figure 4 & 5). This is particularly apparent with the individual ONT libraries assembled with Unicycler which went from obtaining some of the lowest average NGA50 values to having some of the highest (3%, 1% and 0.1% increasing to 43%, 35% and 29% respectively), suggesting the ONT-Unicycler assemblies contained contigs below the 90% ANI threshold required for mapping and were only aligned to the reference genomes post-polishing (Supplementary Figure 4 & 5). A post-polish increase was more modest in PacBio assemblies, which increased from 3.7% to 3.8% and from 4.9% to 5.1% for the Flye and wtdbg2 assemblies respectively (Supplementary Figure 4 & 5). Manual inspection of contig alignments from long-read only assemblies to the reference genomes revealed large numbers of overlapping misassembled contigs that were not resolving into a single assembly. The lack of contiguity is potentially due to the higher error frequency associated with ONT and PacBio reads.

To determine if the long-read assemblies were failing due to high sequencing depth, reads mapping to genomes with ≥100 x coverage were extracted, randomly downsampled to different coverage levels, and re-assembled. For both ONT and PacBio, and all assemblers used, downsampling the reads prior to re-assembly led to more complete assemblies (Figure 4; Supplementary Table 5). Furthermore, successful assemblies using PacBio reads with Unicycler were only obtained after downsampling. However, Nanopore reads assembled with Unicycler obtained the most complete assemblies using the original mixed community reads (i.e., rather than reads mapping to that genome only).

**Figure 4.**
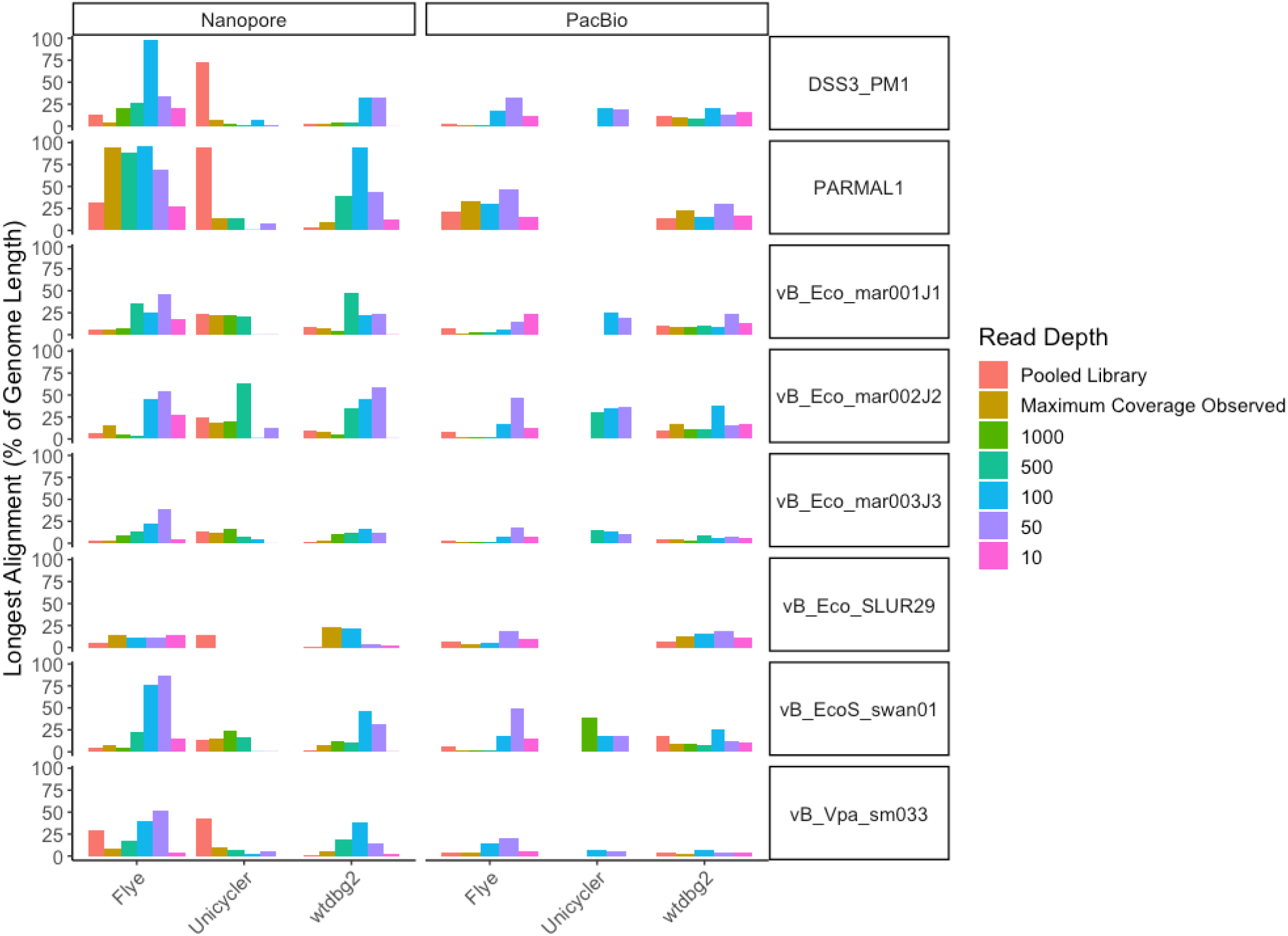
Effect of read depth on long-read assembly. To determine if read depth was a factor in genome assembly when using long reads. Reads were first mapped to individual genomes, then randomly subsampled from the Maximum coverage obtained in a sample, down to give 1000x, 500x, 100x, 50x and 10x coverage per genome, before being assembled. The proportion of the original reference genome (%) that was assembled into a single contig was then determined.

### Assembly Results - SNPs, INDELs and Mis-assemblies

To investigate the fidelity of assemblies, we compared assembled contigs to the mock community reference genomes to identify the frequency of SNPs and INDELs per 100 kb. Assemblies with the highest fidelity would be expected to have the smallest number of SNPs and INDELS compared to the known references. Both SNPs and INDELs were calculated for genomes where ≥ 50% of the genome was covered by contigs. Using Illumina only reads resulted in the lowest number of SNPs per 100 kb (503) with ONT long-read only assemblies having the highest number of SNPs. The number of SNPs in long read assemblies was also dependent on the assembler used. Using ONT reads with Flye (2038) resulted in fewer SNPs than wtdbg2 (3545) or Unicycler (4159) (Supplementary Figure 6A & 7). Conversely, PacBio reads assembled with Flye had a higher SNP frequency (2180) than those produced using wtdbg2 (1806) (Supplementary Figure 6A & 7).

A similar pattern of results was observed for the number of INDELs per 100 kb, although a much larger difference between the different read technologies was observed. Again, the assembler used had an impact on the frequency of INDELs. ONT assemblies produced by far the largest number of INDELS when using Unicycler (miniasm + racon; 4521) compared with Flye (1702) and wtdbg2 (1982) assemblies (Supplementary Figure 6B & 8). PacBio assemblies had far fewer INDELs than ONT with far fewer INDELs observed in Flye assemblies (176) than wtdbg2 assemblies (946). Illumina only assemblies had by far the smallest number of INDELs (14) (Supplementary Figure 6B & 8).

Whilst long-read-only assemblies had a high frequency of SNPs and INDELs, hybrid assemblies produced with Unicycler that combined Illumina reads with ONT or PacBio reads obtained SNP and INDEL levels comparable to Illumina only assemblies (Supplementary Figure 6B). Thus, there is a clear advantage to using a hybrid assembly as it reduces the number of errors.

### Effect of Polishing Long-Read Assemblies on SNPs, INDELs and ORF Prediction

Using short reads to polish contigs produced from long read assemblies generally reduced the number of SNPs per 100 kb, although this was dependent on the specific assembly. Polishing ONT assemblies produced with Unicycler and wtdbg2 decreased the frequency of SNPs by 42% and 26%, respectively (Figure 5A). Polishing the Flye assembly resulted in a small increase in the number of SNPs (Figure 5A). Rather than introducing errors, this is likely as a result of contigs prior to polishing having SNP frequencies that prevented recruitment to a reference genome at 90% identity by mapping. Post polishing, these contigs are now recruited to genomes, but still contain a number of SNPs (Figure 5A). With PacBio reads assembled with Flye, polishing had no effect on the number of SNPs (Figure 5A). For the PacBio wtdbg2 assembly, the number of SNPs increased, as observed with ONT reads assembled with Flye. Again, this increase is likely due to the increased number of contigs that are mapped to the reference genome (Figure 5A).

**Figure 5.**
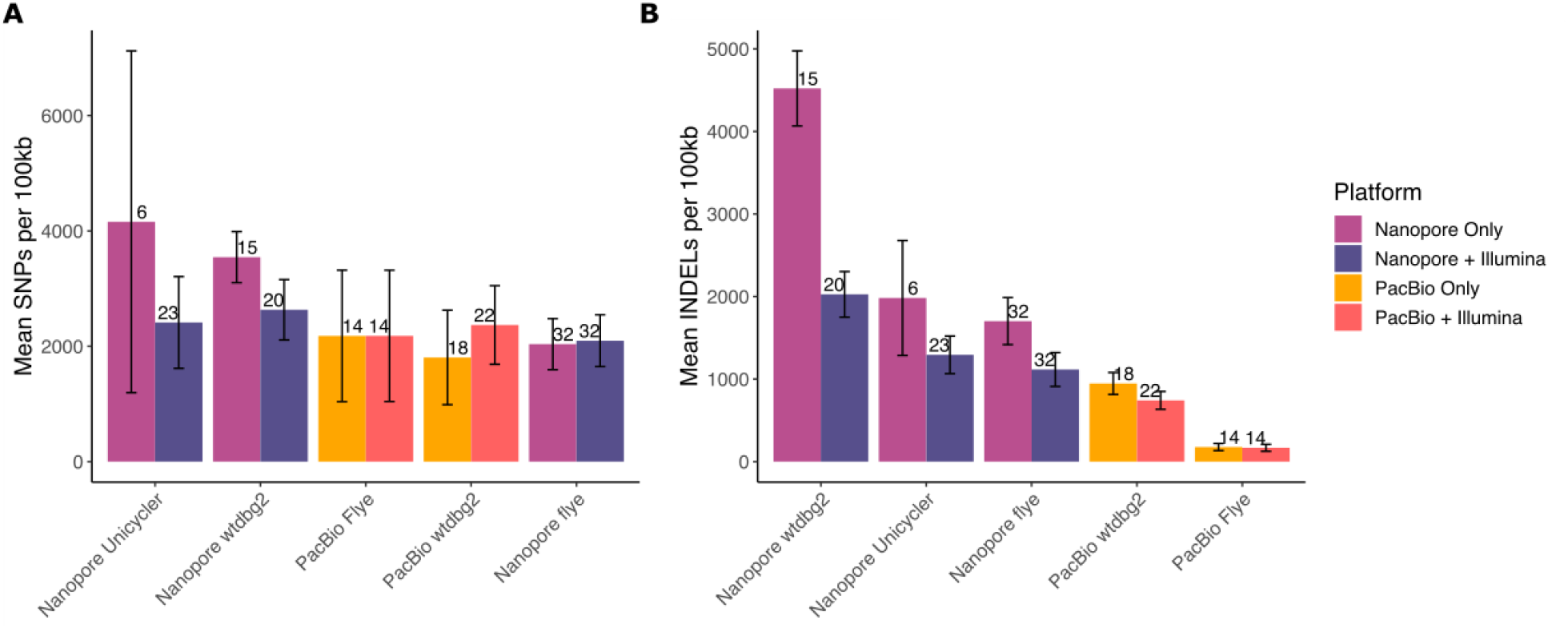
Effect of polishing on error rate. SNPs and INDELs were identified in contigs from nanopore only, nanopore contigs polished with illumina reads, PacBio only and PacBio contigs polished with illumina reads. The number of SNPs (A) and INDELSs are expressed as the number per 100 kb of reference genome, where at least 50% of the reference genome was recovered by contigs, where at least 50% of the reference genome was recovered by contigs. Error bars are standard error of the mean, and the number above the bar indicates the number of genomes included in the mean calculation (from a total possible maximum of 60 (15 genomes, 4 assemblies)).

The effect of polishing on the frequency of INDELs was more apparent. The ONT assemblies had a higher number of INDELs than PacBio assemblies prior to polishing (Figure 5B) and thus were of lower quality. For ONT reads assembled with Unicycler (miniasm + racon), which had the highest frequency of INDELs initially, there was a 55% decrease in INDELs post polishing (Figure 5B). For ONT reads assembled with wtdbg2 and Flye, there was a ~34% decrease in the number of INDELs per 100 kb (Figure 5B). For PacBio assemblies the starting frequency of INDELs was lower than ONT prior to polishing but polishing with Illumina reads still resulted in a 21% and 4.5% decrease in INDEL frequency for wtdbg2 and Flye assemblies respectively (Figure 5B).

As assembly errors can have an effect on ORF prediction and functional annotation [58], we investigated the number and length of predicted ORFs on contigs which mapped to reference genomes before and after polishing. Polishing with short reads had the greatest effect on ONT data regardless of the assembler used, with mean ORF length increasing for all assemblies. Both Unicycler and wtdgb2 observed mean ORF length increases of ~66%, with a ~24% increase for Flye (Figure 6). For PacBio assemblies, the increases in mean ORF length were more modest at ~11% for wtgb2 assemblies and ~0.2% for Flye assemblies (Figure 6). While there was an increase in mean ORF length for all combinations of reads and assemblers post-polishing, all combinations were still smaller than the value obtained for the 15 reference genomes (Figure 6).

**Figure 6.**
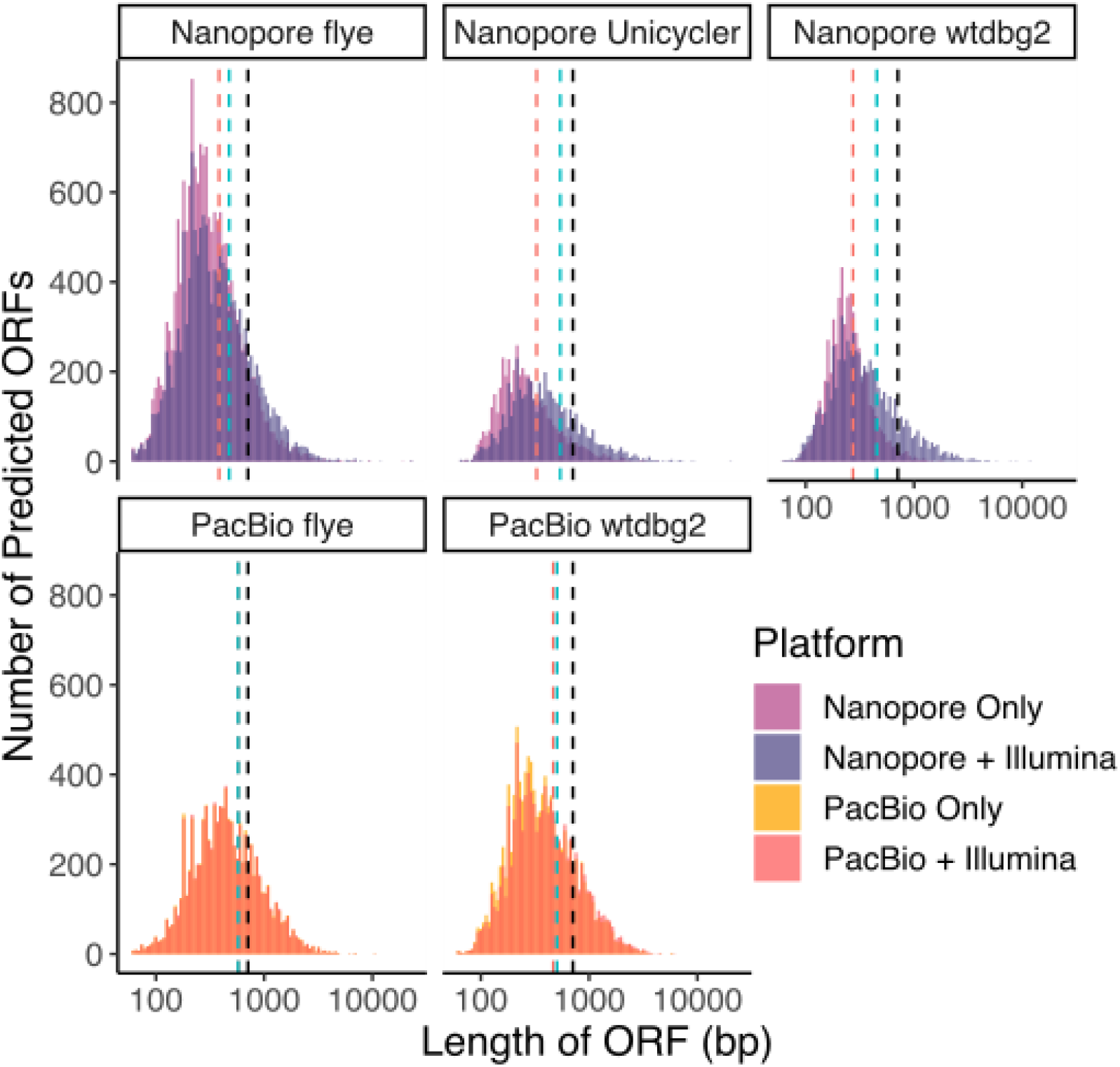
The effect of polishing long-read assemblies on predicted ORF lengths. ORF length was calculated from nanopore only contigs, nanopore contigs polished with illumina reads, PacBio only contigs and PacBio contigs polished with illumina reads. With contigs obtained using Flye, wtdbg2 and Unicycler assemblers. Histograms show the distribution of predicted ORF length for sequencing type (Nanopore or PacBio) and assembly algorithm. The expected mean ORF length from the reference genomes is represented as dashed vertical line (709 bp; black), compared to the mean value before (red) and after (blue) polishing.

### Effect of Polishing Long-Read Assemblies on Viral Prediction

Many viral prediction programs use similarity of predicted proteins to known hallmark proteins for virus prediction. Thus, truncated proteins may alter the ability to predict viral contigs from viromes. To test if truncated proteins affect virus prediction, we compared VIBRANT [47] which in part uses predicted proteins, and DeepVirFinder [48] a K-mer based prediction system on all assembled contigs. Although we utilised purified phage isolates to create the mock community, up to 20% of the reads from Illumina libraries did not map to the reference genomes. Therefore, we utilised this unfortunate level of contaminating host bacterial DNA for benchmarking viral prediction. To determine how many predictions represented “true” viral predictions, we mapped the predicted vOTUs against the reference genomes.

For DeepVirFinder predictions, there were minimal differences in the number of predicted viral contigs (vOTUs) before and after polishing for all assemblies. The largest difference was observed for ONT reads assembled using Flye (61 before, 52 after; Supplementary Figure 9; Supplementary Table 7). However, there was a marked increase in the number of vOTUs that could be verified as phage. For Flye, the number that could be verified as phage increased from 82% to 96% after polishing, wtdbg2 assemblies increased from 83% to 98%, and Unicycler assemblies increased from 93% to 99%. Thus, polishing ONT assemblies with Illumina reads led to an overall decrease in the number of erroneous viral predictions when using DeepVirFinder (Supplementary Figure 9; Supplementary Table 7). For the PacBio assemblies, there was no difference in the number of predicted vOTUs and those that could be verified as phage when using DeepVirFinder (Supplementary Figure 9; Supplementary Table 7).

When using VIBRANT for prediction, polishing of PacBio assemblies had no or minimal effect on the number of predictions or the number of verified predictions (Supplementary Figure 9; Supplementary Table 7). However, the polishing of ONT assemblies led to vastly different numbers of predicted vOTUs, and this varied with <u>the</u> assembler used. The largest difference was for the ONT wtdbg2 assembly, decreasing from 199 to 133 predicted vOTUs, and the proportion of verified phages increased for all ONT assemblies after polishing. For Flye, the number of verified phages increased from 75% to 81%, Unicycler increased from 72% to 96%, and wtdbg2 increased from 51% to 87% (Supplementary Figure 9; Supplementary Table 7).

Thus, when using DeepVirFinder there was minimal impact of polishing on the prediction of vOTUs from either PacBio or ONT assemblies. However, there were clear benefits to the polishing of ONT assemblies when using VIBRANT for vOTU prediction, as the percentage of vOTUs that could be verified to be phage increased post polishing.

### Effect of sequencing technology on predicted virome diversity

Having established DeepVirFinder generally performed better for all sequencing technologies, we utilised the output of DeepVirFinder predictions to assess how diversity statistics of the mock community varied with sequencing technology and assembly.

Alpha diversity was assessed using predicted vOTUs, Shannons’s index and Simpson’s index; overall the same general trend was observed for each metric (Figure 7; Supplementary Table 8). When using long read assemblies there was an overprediction of alpha diversity (ONT: median vOTUs 54.5, PacBio: median vOTUs 77.5; Known number vOTUs 15). In contrast, when using Illumina and Illumina + ONT/PacBio hybrid there was an underestimation of alpha diversity (ONT-Illumina hybrid: median vOTUs 8.5, PacBio-Illumina hybrid: median vOTUs 6.5, Illumina: median vOTUs 6.5). Within this general trend, there was further variation with the method used for assembly. While the use of Illumina or Illumina +ONT/PacBio generally led to an underestimation of diversity, it was closer to the known diversity measurement than the use of long reads alone. When using long reads alone, using ONT assembled with Unicycler give the closest estimate of the true diversity of the mock community.

**Figure 7.**
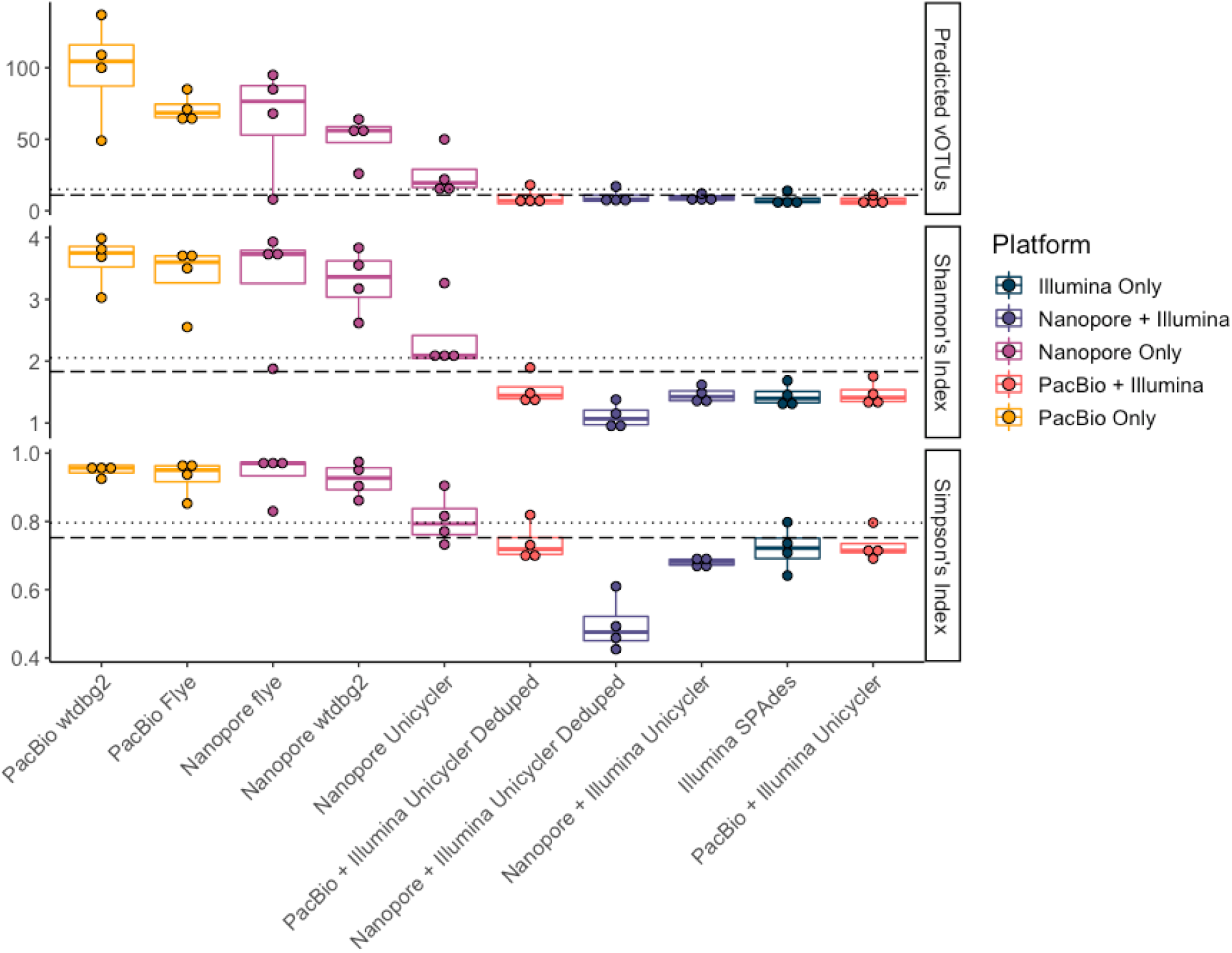
The effect of sequencing platform and assembler on diversity estimates. Boxplots showing the number of predicted vOTUs for mock virome analysis (top), and Shannon’s index (middle) and Simpson’s index (bottom) alpha diversity measures. Dotted lines indicate true values for mock virome input, and dashed lines indicate true values excluding genomes that were not detected by read mapping in any library.

## Discussion

The use of long read sequencing technologies is becoming increasingly common for the sequencing of metagenomics samples, in particular those that focus on the bacterial community. A number of studies have demonstrated the advantage of long-reads in assembling complete genomes from a variety of samples [25, 59–61]. There have also been a number of studies benchmarking the assembly and/or recovery of bacteria from mock communities using long-reads [62, 63], along with benchmarking of assembly algorithms for prokaryotic genomes (excluding phages) [64, 65]. However, there are no such comprehensive studies that have directly compared Illumina, ONT, and PacBio sequencing technologies for the study of viromes.

Previous benchmarking of short-read assemblers has demonstrated minimal differences in genome recovery of phage genomes when comparing multiple assemblers on a mock viral community [51]. For this reason, we chose only one short-read assembly algorithm: SPAdes. For long-read assembly, we chose three frequently used approaches of Unicycler (miniasm + racon), Flye, and wtdgb2 as well as using Unicycler for a direct hybrid assembly. For long read sequencing alone, we were unable to obtain assemblies from PacBio reads with Unicycler, even when combining all three libraries, suggesting it was not a lack of coverage.

When using a single sequencing technology, only Illumina reads resulted in the complete assembly of a phage genome within any sample. Utilising a hybrid approach increased the number of genomes that could be assembled, with ONT + Illumina reads assembled with Unicycler (miniasm + racon) recovering the largest number of genomes, whereas the addition of PacBio reads did not result in the same increased recovery of genomes. However, this may well be due to the reduced yield of PacBio reads compared to ONT reads. Thus, increased yield of PacBio data might improve this metric. However, it was clear for both ONT and PacBio that very high coverage of specific genomes within a sample was detrimental to genome assembly, with subsampling of reads improving the length of the contig recovery. Thus, digital normalisation of reads may improve recovery of longer contigs as has been found in other systems [66]

The combination of long and short reads improving recovery of assembled genomes is consistent with previous benchmarking of a mock viral community using a virION approach [14]. Unlike the virION approach, we were only able to assemble a complete genome with just long-reads after downsampling to lower read depths prior to assembly. However, direct comparison between the studies is difficult given the different phages used in each mock community. Here, we utilised MDA application to provide sufficient material for long-read sequencing, whereas the virION utilises PCR to provide sufficient material [14, 23]. The virION approach has comprehensively demonstrated that the relative abundance of phages is maintained due to the LASL-PCR approach [14, 23]. Here, we observed a strong correlation in the abundance of phages in the un-amplified Illumina viromes and amplified long-read viromes. However, we are cautious in the interpretation of this data. The DNA from a ssDNA phage (ΦX174) was spiked into our mock community at a deliberately low level, as we wanted to avoid flooding our amplified DNA with ssDNA given known biases of MDA. However, given the lack of detection of ΦX174 in any samples, we may have been overly cautious in the amount added. Thus, when ssDNA phages are present in a community, it is likely the biases observed previously are still likely to hold true [19–21].

When assessing any individual sequencing technology alone, the lowest number of SNPs or INDELs observed when using Illumina reads, with ONT assemblies having a larger number of SNPs, and in particular INDELs, compared to PacBio assemblies. Thus, the short read assembly produced the highest fidelity genomes. Both INDELs and SNPs were also affected by the method used for assembly. For ONT reads, Flye produced assemblies with the lowest number of INDELs or SNPs compared to wtdgb2 and Unicycler (miniasm + racon). It is likely for ONT data that the number of SNPs and INDELs will further decrease with improvements in accuracy reported for both R10 flow cells and the latest base calling algorithms that have been developed since this data was collected, as this data was generated with R9 flow cells. In contrast, Flye assemblies of PacBio reads had the lowest number of SNPs, but the highest number of INDELs. Thus, the choice of assembly method should be adjusted for the type of long-reads being used. The addition of short reads to polish the long read assemblies resulted in a reduction of both SNPs and INDELs, as has been observed in other studies [14, 22, 26].

While the combination of both short Illumina reads with long reads resulted in the “best” overall assemblies, it may well not be feasible to sequence samples with both technologies. Therefore, we treated the assemblies from multiple approaches to assess how the different approaches affected the predicted diversity of the sample. Although polishing long read assemblies had a significant impact on reducing the number of SNPs and INDELs, there was minimal effect on the number of predicted contigs that were viral when using DeepVirFinder for prediction. However, VIBRANT, which in part utilises the identification of hall-mark phage genes and was more sensitive to the higher error rates of un-polished long-read assemblies, obtained far fewer erroneous viral predictions post-polishing. Thus, choice of sequencing technology may have ramifications for downstream choices in viral prediction software.

## Conclusions

We have benchmarked Illumina, ONT, and PacBio sequencing platforms for virome analysis using a number of read and assembler combinations and offer recommendations for the community: (i) if only using one sequencing platform, Illumina performs best at genome recovery and has the lowest error rates; (ii) the addition of long-reads to Illumina reads improves the assembly of lowly abundant genomes, particularly ONT; (iii) whilst long read assemblies, particularly ONT, have higher error frequencies, polishing with Illumina reads can reduce these errors to levels comparable with Illumina-only assemblies; (iv) down-sampling of long reads may aid assembly; and (v) the choice of sequencing platform should be considered when making downstream analysis decisions, such as assembler algorithm and viral prediction software.

## Abbreviations

INDEL: Insertion or Deletion
LASL: Linker Amplified Shotgun Library
MDA: Multiple Displacement Amplification
ONT: Oxford Nanopore Technology
SNP: Single Nucleotide Polymorphism
TFF: Tangential Flow Filtration
vOTU: Viral Operational Taxonomic Unit

## Data Availability

All reads from virome sequencing were submitted to the ENA under study PRJEB56639. The assemblies are provided via FigShare (10.25392/leicester.data.21346935)

## Acknowledgments

Some text

## Author Contributions

NB &, DS, AM conceived the idea. AM, BR, SM, TR, AN, YC, MC and DJS provided funding and/or strains. AM, DJS, DS, MJ and JH supervised the work. RC analysed the data. RC and AM drafted the manuscript, with all authors contributing to editing of the final manuscript.

